# ESCRT-0 is not required for ectopic Notch activation and tumor suppression in *Drosophila*

**DOI:** 10.1101/002790

**Authors:** Emiliana Tognon, Nadine Wollscheid, Katia Cortese, Carlo Tacchetti, Thomas Vaccari

**Affiliations:** IFOM, Istituto FIRC di Oncologia Molecolare @ IFOM-IEO Campus via Adamello 16, 20139, Milano, Italy; DIMES, Dipartimento di Medicina Sperimentale, Anatomia Umana, Università di Genova, Via de Toni 14, 16132, Genova, Italy; Experimental Imaging Center, Scientific Institute San Raffaele,Via Olgettina 60, 20132, Milano, Italy

**Keywords:** ESCRT-0, Hrs, Stam, JAK/STAT signaling, Notch Signaling, Endocytosis

## Abstract

Multivesicular endosome (MVE) sorting depends on proteins of the Endosomal Sorting Complex Required for Transport (ESCRT) family. These are organized in four complexes (ESCRT-0, -I, -II, -III) that act in a sequential fashion to deliver ubiquitylated cargoes into the internal luminal vesicles (ILVs) of the MVE. *Drosophila* genes encoding ESCRT-I, -II, -III components function in sorting signaling receptors, including Notch and the JAK/STAT signaling receptor Domeless. Loss of ESCRT-I, -II, -III in *Drosophila* epithelia causes altered signaling and cell polarity, suggesting that ESCRTs genes are tumor suppressors. However, the nature of the tumor suppressive function of ESCRTs, and whether tumor suppression is linked to receptor sorting is unclear. Unexpectedly, a null mutant in *Hrs,* encoding one of components of the ESCRT-0 complex, which acts upstream of ESCRT-I, -II, -III in MVE sorting is dispensable for tumor suppression. Here, we report that two *Drosophila* epithelia lacking activity of *Stam*, the other known components of the ESCRT-0 complex, or of both *Hrs* and *Stam* fail to degrade signaling receptors. However, mutant tissue surprisingly maintains normal apico-basal polarity and proliferation control and does not display ectopic Notch signaling activation, unlike cells that lack ESCRT-I, -II, -III activity. Overall, our *in vivo* data indicate that the ESCRT-0 complex plays no crucial role in regulation of tumor suppression, and suggest re-evaluation of the relationship of signaling modulation in endosomes and tumorigenesis.

## Introduction

Epithelial tissue development and homeostasis relies on proper coordination of cell polarity and cell growth. Cell-cell communication enables such coordination via a number of conserved signaling pathways. Consistent with this, deregulation of signal transduction frequently alters cell polarity and growth and is commonly observed in pathology.

A major modulator of signaling outputs is endocytic trafficking [1]. Underscoring the importance of endocytosis in modulation of a number of signaling pathways, endocytic proteins are increasingly found mutated in cancer [2]. In most pathways, initiation of the signaling cascade occurs at the plasma membrane, when ligands meet their cognate receptors. Subsequent internalization of ligand-receptor cargo complexes usually leads to transport to early endosomes. Following endosomal entry, receptors can be recycled back to the plasma membrane for further rounds of signaling, or destined degradation in the lysosome. Both fates can potentiate or attenuate signaling depending on the specific mechanisms of signaling activation of each receptor and on the handling of other signaling components by the endocytic machinery. For example, while some receptors continue to signal in endosomes, as is the case of some Receptor Tyrosine Kineses (RTKs), others require recycling back to the plasma membrane, such as the Transferrin receptor [1].

Endosomal sorting is the entry point into the degradative fate and it involves sorting of ubiquitylated cargoes on the limiting membrane of endosomes and the formation of Multi Vesicular Endosomes (MVEs). Endosomal sorting and MVE biogenesis are controlled by Endosomal Sorting Required for Transport (ESCRT) proteins. Four multi-subunit ESCRT complexes (ESCRT-0, -I, -II, -III) act in a sequential fashion to deliver cargoes into the internal luminal vesicles (ILVs) of the nascent MVE [3–5]. The process is thought to start when the ESCRT-0 components Hrs and Stam, acting as an heterodimer, clusters ubiquitylated cargoes in flat clathrin-coated domains of the endosomal membrane. ESCRT-0 then is thought to recruit the ESCRT-I complex and subsequent action of ESCRT-II and -III complexes leads to de-ubiquitylation of cargoes and their sequestration in forming ILVs [6–9]. The full extent of cargoes subjected to endosomal sorting, and how sorting affects signaling modulation precisely is largely unknown.

Mutants for ESCRT components in metazoan animals, such as *Drosophila melanogaster,* have been recently providing a fascinating initial glimpse in the importance of endosomal degradation for signaling regulation during development [10]. In fact, in addition to showing failure to degrade a number of transmembrane signaling receptors, they show ectopic activity of multiple signaling pathways, including Notch, JAK/STAT and others [11–16]. In addition, epithelial tissue mutant for a large number of ESCRT genes display altered apico-basal polarity and unrestrained proliferation leading to formation of tumor-like masses, indicating that endosomal sorting, possibly by regulating signal transduction play a major role in tumor suppression [17, 18]. Loss of tumor suppression in *Drosophila* ESCRT mutants requires ectopic activity of Notch, JAK/STAT, and dpp and JNK signaling, as down-modulation of these pathways in ESCRT mutant rescues the overproliferation or the loss of polarity, or both. For instance, ESCRT mutant cells display (and rely on) ectopic, ligand-independent activation of Notch signaling for cell-autonomous proliferation and on ectopic JAK/STAT signaling activation for cell-autonomous non cell-autonomous proliferation [11]. Such dramatic increase of proliferative signaling alters cell cycle regulation and is counteracted by JNK-and Hippo-mediated mediated activation of apoptosis [14, 16]. Thus, while the proliferative defects of ESCRT mutants are well documented, how apico-basal polarity is compromised is still obscure. Despite this, consistent with conservation in the involvement of ESCRTs in tumor suppression, a number of ESCRT-I, -II, -III components have been found mis-expressed in various cancers [see for review 19].

Unexpectedly, while all the *Drosophila* ESCRT-I, -II, -III genes analyzed so far behave as tumor suppressors and prevent ectopic ligand independent Notch activation, Drosophila *Hrs,* which encodes for one of the two obligate ESCRT-0 components, is required for endosomal sorting, and signaling attenuation by RTKs, but it appears dispensable for tumor suppression. In addition, in a *Hrs* mutant, Notch fails to be degraded but it is otherwise normally activated [20–22]. It has been recently reported that mutants in *Stam,* which encodes for the Hrs partner in ESCRT-0, and *Hrs Stam* double mutants affect endosomal sorting, MVE biogenesis and alter RTK signaling [23, 24]. However, it has not been tested whether *Stam* or *Hrs Stam* double mutants display loss of tumor suppression or altered Notch trafficking and signaling. Thus, we decided to analyze epithelial tissues that lack function of Stam or both Hrs and Stam during *Drosophila* development.

Here we show that differently from ESCRT-I, -II, -III mutants, *Stam* or *Hrs, Stam* double mutants do not present loss of tumor suppression or ectopically active Notch signaling. However, similarly to single *Hrs* mutants and other ESCRT mutants, *Stam* or *Hrs, Stam* double mutants display endosomal accumulation of ubiquitinated cargoes, including Notch and the JAK/STAT receptor Domeless. Unexpectedly, our data indicate that ESCRT-0 is dispensable for tumor suppression and ectopic Notch signaling activation, and shed light on the mechanism of ESCRT-mediated tumor suppression and of endosomal Notch activation.

## Materials and Methods

### Fly strains and genetics

*Drosophila* lines referred to in the text are *Hrs^D28^* [20], *Stam^2L2896^* [24], and the double mutant *Hrs^D28^ Stam^2L2896^* (Bloomington Drosophila Stock Center (BDSC) #3914, #41804 and #41806, respectively). Predominantly mutant eye and wing discs (referred to in the text as mutant discs) were generated with the eyeFLP cell lethal system as described [25]. Mutant eye disc clones were generated with the eyeFLP mosaic system as described previously [26]. Mutant FE cell clones were generated by using the heat shock-mosaic system [27] and the GR1 system [28]. For most of the mosaic experiments, female flies were heat-shocked at 37 °C for 1 h two times a day for 2 days and then incubated at 25 °C for 4 days before dissection. Detailed genotypes are available upon request.

The *Hrs*, *Stam* recombinants devoid of *l(2)gl* lesions were generated via standard genetic procedures. After we made sure that both the *Hrs^D28^* and *Stam^2L2896^* single mutants did not contain *l(2)gl* lesions by complemention assay with the null allele *l(2)gl^4^*, *Hrs^D28^* females were crossed with *Stam^2L2896^* males to generate recombinogenic F1 females. These were then crossed to a balancer stock and the F2 male progeny was stocked and crossed back to *Hrs* and *Stam* mutants and relative deficiencies (*Hrs* deficiency: BDSC #9543; *Stam* deficiency BDSC #7821). Males that failed complementation with both loci but complemented *l(2)gl^4^* or a *l(2)gl* deficiency (BDSC #3634) were kept as independent recombinant fly lines.

### Immunostainings and confocal microscopy

Ovaries and discs were dissected in S2 medium (Invitrogen) containing 10% fetal bovine serum and PBS respectively, fixed in 4% formaldehyde for 10 minutes at room temperature and then rinsed three times in phosphate buffered saline with 0.3% Triton X-100 and 0.5% BSA. Primary antibodies were used for immunostaining against the following antigens: Hnt, Cut, Notch ECD, Notch ICD, (all from Developmental Studies Hybridoma Bank- DSHB); Dome (A gift from Stephane Noselli). Avl (Lu and Bilder, 2005); Ubiquitin FK2 (Biomol); activated Caspase-3 (Signal Transduction Technologies). Secondary antibodies conjugated to Alexa-488, Alex-568 were used (Molecular Probes). Phallodin-TRITC from sigma was used to mark F-actin while DAPI (4’6-diamidino-2-phenylindole) to stain the nuclei. The images were obtained using a Zeiss LSM 510-Meta confocal microscope or aa TCS microscope (Leica). Images were edited with Adobe Photoshop CS and were assembled with Adobe Illustrator.

### Trasmission electron microscopy

Eye discs WT or mutant for *Stam*, *Hrs, Stam l(2)gl* or *Vps25* were fixed in 2.5% glutaraldeyde diluted in 0.1 M sodium cacodylate buffer for 3 hours at room temperature. Eye discs were post-fixed in 1% osmium tetroxide (Electron Microscopy Science, Hatfield, PA, USA) for 2 hours at room temperature and subsequently in 1% uranyl acetate (Electron microscopy science) for 1 hour. Samples were dehydrated through a graded ethanol series and next in propylene oxide before embedding in epoxy resin (Poly-Bed, Polyscience, Warrington, PA, USA) overnight at 42°C and then 2 days at 60°C. Searching for the eye disc epithelium was performed on semi-thin sections (500 nm) stained with toluidine blue. Ultrathin sections of 50 nm were then cut and stained with 5% uranyl acetate and lead citrate. Representative TEM micrographs of each sample were taken with Tecnai 12-G2 microscope (FEI company, Eindhoven, The Netherlands) and processed with Adobe Illustrator CS5.

### Notch-trafficking Assay

Wild-type or eyFLP/+; FRT40A *Stam^2L2896^*/FRT40A P(mini-w, cl), eyFLP/+; *FRT40A Hrs^D28^ Stam^2L2896^ l(2)gl* FRT40A P(mini-w, cl) eye discs were dissected in Schneider’s *Drosophila* medium and after dissection the medium was replaced. Imaginal discs were cultured for 20 and 60 min, respectively, in presence of anti-Notch ECD antibody that recognizes the extracellular portion of Notch. Following medium changes the organs were fixed immediately for the 0 min time point or after 60 min or 300 min for the different time points. Localization of the anti-Notch EDC antibodies was revealed using secondary antibody, and co-staining with anti-Avl was performed in a subset of samples.

### RT-PCR

Total RNA from wing imaginal discs (40 discs per sample) was extracted using TRIZOL Reagent (Invitrogen) and RNeasy Mini Kit (Qiagen) according to the manufacturer’s protocol. Concentration and purity was determined by measuring optical density at 260 and 280 nm using a Nanodrop spectrophotometer. 1 μg of total RNA was reverse transcribed using a SuperScript VILO cDNA Synthesis kit (Invitrogen) according to the manufacturer’s protocol. 5 ng of cDNA was amplified (in triplicate) in a reaction volume of 15 μl containing the following reagents: 7.5 μl of TaqMan PCR Mastermix 2x No UNG (Applied Biosystems, Foster City, CA), 0.75 μl of TaqMan Gene expression assay 20x (Applied Biosystems, Foster City, CA). For each sampls 300 nM of primers and 100 nM of Roche probes were used. RT-PCR was carried out on the ABI/Prism 7900 HT Sequence Detector System (Applied Biosystems), using a pre-PCR step of 10 min at 95°C, followed by 40 cycles of 15 s at 95°C and 60 s at 60°C. The following primers were used: Hrs: fwd tcaaccagaaagatgtcactcc; rev ccaggagggaatagcagga; Stam: fwd ggaatctttgggcagtcgt; rev ccagttgtcgttggtattagtttc; Vps25: fwd ccttcccacccttctttaca; rev tgcctgaggtatttgagaaagag; RpL32-RA: fwd cggatcgatatgctaagctgt; rev cgacgcactctgttgtcg;

## Results

### The reported ESCRT-0 double mutant allele contains a *l(2)gl* mutation

To compare the phenotype of the *Stam* or of the *Hrs, Stam* double mutant to that of *Hrs* or of ESCRT-I, -II, -III mutants, we generated clones of cells mutant for *Stam^2L2896^* (Mutant cells are GFP-negative; see Material and Methods) in the follicular epithelium (FE) of the Drosophila ovary. As it is the case of FE cells mutant for *Hrs^D28^*, *Stam* mutant FE cells display normal epithelial morphology (Fig. 1A–C). In contrast, cells homozygous for a recently reported *Hrs^D28^, Stam^2L2896^* double mutant allele [23] formed large clones of mesenchymal-like cells (Fig. 5A). We observed a similar phenotype when we generated mosaic eye imaginal discs or eye imaginal discs consisting predominantly of cells mutant for *Hrs*, *Stam* or both (Fig. 1E–G, M–O; Fig. 5B, D). Both *Hrs* and *Stam* are on chromosome 2L, which harbors in a sub-telomeric position the tumor suppressor *l(2)gl*, a gene frequently lost by spontaneous deletion [29]. Thus, we wondered whether the *Hrs, Stam* double mutant chromosome carried a mutation in *l(2)gl.* Failure to complement the null allele *l(2)gl^4^* indicated a possible lesion in *l(2)gl* on the chromosome carrying both *Hrs* and *Stam* mutations. To test if it was indeed the case, we recombined away the distal part of chromosome 2L containing *l(2)gl* from the *Hrs Stam* chromosome and retested for complementation. We isolated several independent recombinants that fail to complement *Hrs* and *Stam* deficiencies but complement *l(2)gl^4^,* a further indication of the presence of *l(2)gl* mutation in the original *Hrs Stam* chromosome (*Hrs Stam l(2)gl* triple mutant henceforth; see Material and Methods)

**Figure 1.**
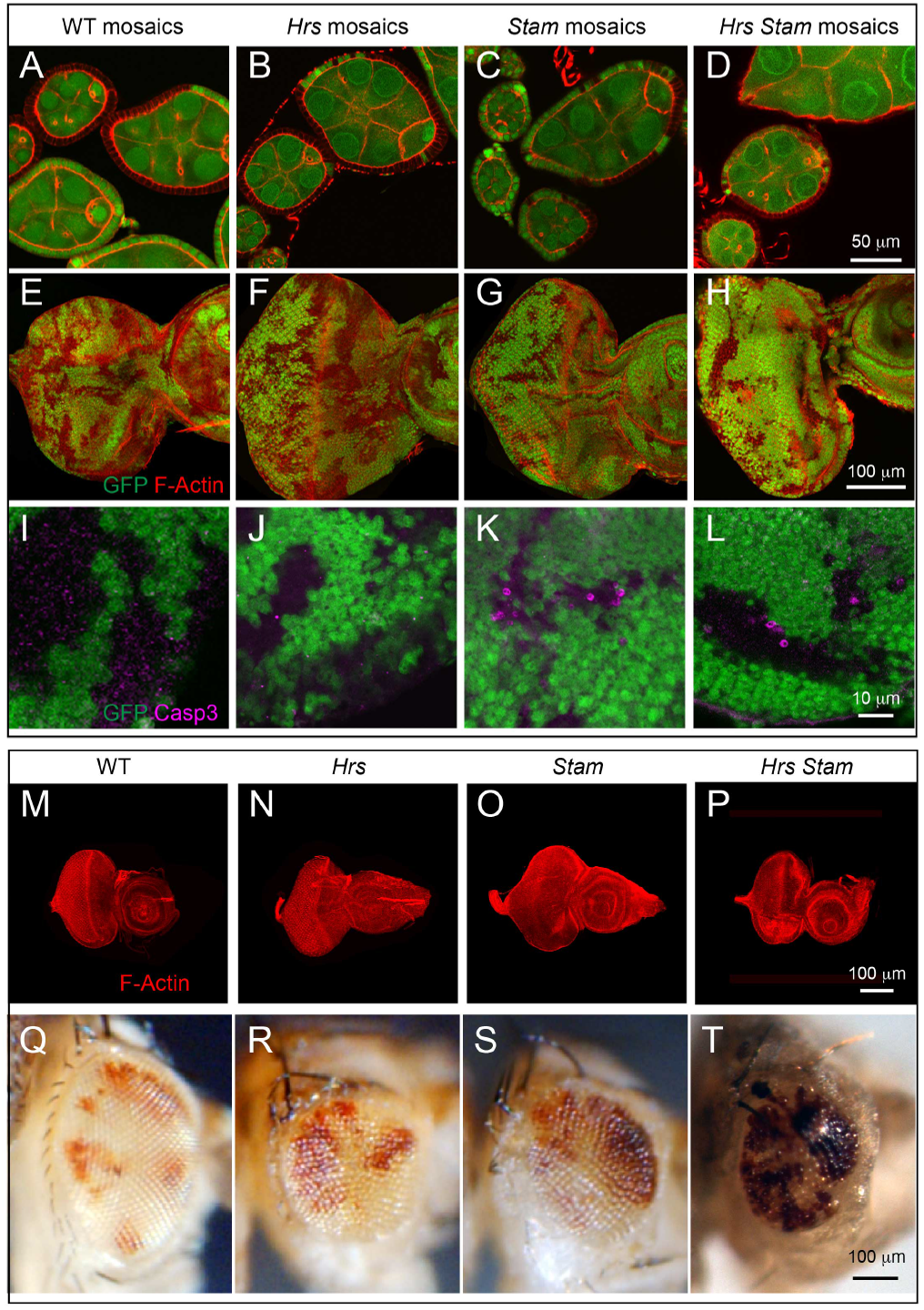
*Hrs*, *Stam* or *Hrs*, *Stam* double mutant tissue do not display altered tissue architecture. (A-H) Epithelial morphology of mosaic FE cells (A-D) and eye discs (E-H) revealed by phalloidin staining to detect F-actin. Follicle cells of 5-7 stage egg chambers homozygous for the mutations (GFP-negative) show normal epithelial architecture compared to WT (GFP-positive). Eye disc cells homozygous for the mutations (GFP-negative) do not show any disruption of tissue architecture.
(I-L) High magnification of a region of mosaic eye imaginal discs. Homozygous cells are marked by the absence of GFP. Apoptotic Caspase-3 (magenta) is activated cell autonomously in a subset of *Hrs* and *Stam* as well as *Hrs*, *Stam* mutant cells, compared to WT.
(M-P) WT and predominantly mutant eye-antennal discs for the indicated gene stained with phalloidin revealed that *Hrs*, *Stam* mutant discs form morphologically normal eye-antennal discs.
(Q-T) Adult eyes deriving from mosaic discs of the indicated genotype. Clones or WT (Q) or mutant cells (R-T) are marked by the absence of red pigment in bright field images indicating that mutant tissue can form photoreceptors.

### ESCRT-0 components are not required for tumor suppression in Drosophila

To test whether the *Hrs^D28^, Stam^2L2896^* mutant chromosome devoid of the *l(2)gl* mutation still possessed tumor-promoting ability, we analyzed mosaic FE, mosaic eye discs, or eye discs consisting predominantly of cells mutant for the recombined allele. Interestingly, these do not display loss of tissue architecture (Fig. 1D, H, P), as is the case of single *Hrs* or *Stam* mutant alleles, suggesting that the *l(2)gl* lesion in the original double mutant allele was responsible for the loss of tumor suppression phenotypes. These data indicate that simultaneous loss of both ESCRT-0 components do not lead to loss of tissue architecture, a striking difference to ESCRT-I, -II, -III mutations, which are tumorigenic [11, 17, 18]. Consistent with this surprising difference, we found that eye discs consisting predominantly of cells mutant for *Hrs*, or *Stam* or both *Hrs* and *Stam* progress to form adult eyes. These are smaller than wild-type and have a rough appearance but contain some mutant photoreceptors (Fig. 1Q–T) The scarcity of mutant adult photoreceptors might be due to cell death, as we occasionally see apoptotic cells in clones of *Hrs*, or *Stam* or both *Hrs* and *Stam* double mutants (Fig. 1I–L). In sheer contrast to these, a number of ESCRT-I, -II, -III mutations, such as those mapping to *Tsg101, vps28, Vps25, vps20,* when made homozygous in eye discs, display a Mutant Eye No Eclosion (MENE) phenotype that have been associated loss of tumor suppression in Drosophila [30]. Overall, these data suggest that the activity of *Hrs* and *Stam* is not tumor suppressive in two different Drosophila epithelial tissues.

### ESCRT-0 components are required for degradation of Ubiquitylated cargoes

To test whether ESCRT-0 mutants are able to sort ubiquitylated cargoes, we immunostained mosaic eye disc and FE cells containing clones of cells mutant for *Hrs,* or *Stam* or both *Hrs* and *Stam* with an antibody specific to mono- and poly-ubiquitin chains. In contrast to WT cells, but similarly to previous reports of *Hrs* and of ESCRT-I, -II, -III mutants [20–22], *Hrs, Stam* and *Hrs Stam* mutant cells, as well as *Hrs Stam l(2)gl* triple mutant cells accumulated ubiquitin (Fig. 2A–F; Fig. 5E–F).

**Figure 2.**
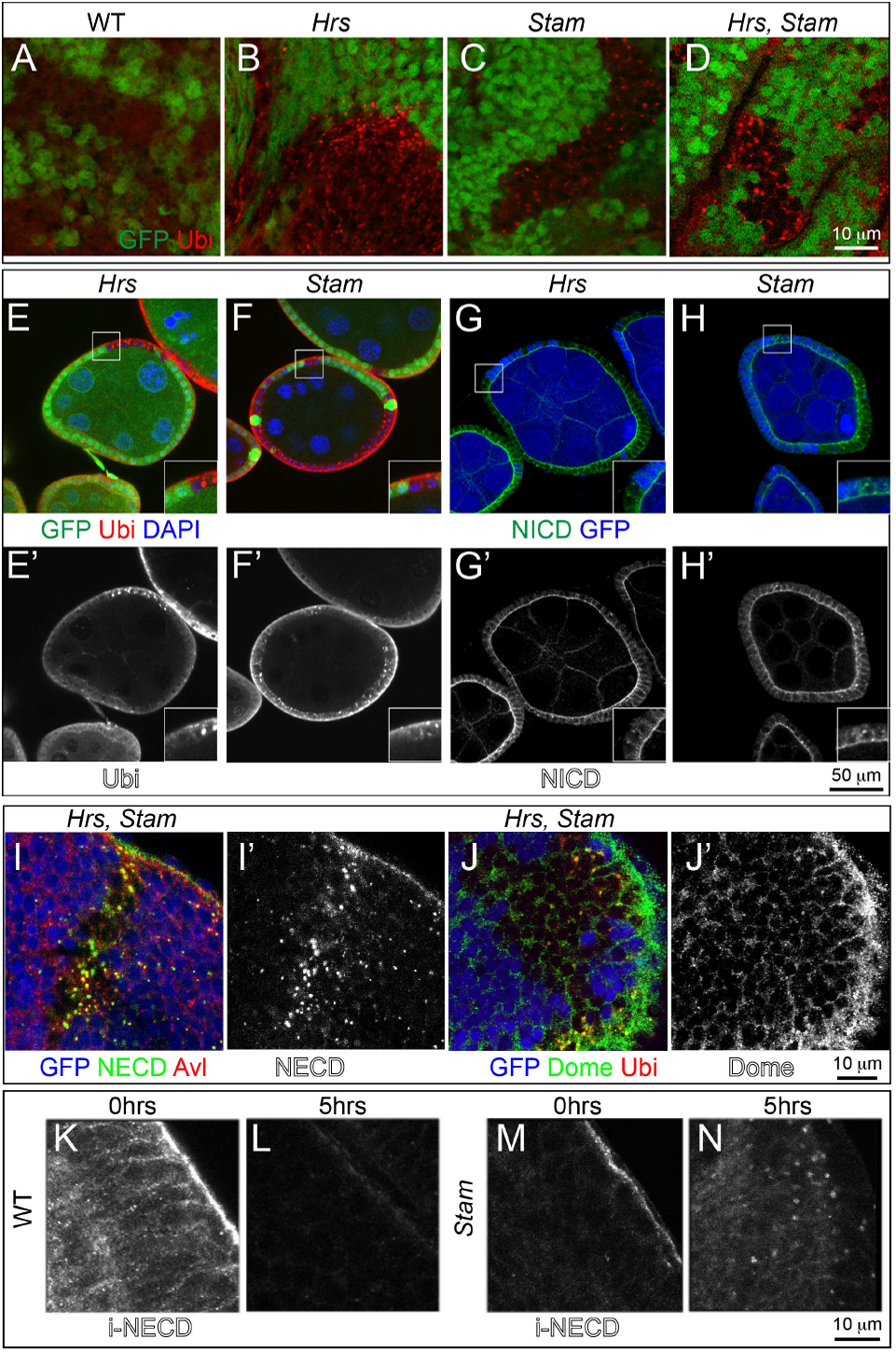
ESCRT-0 mutations lead to accumulation of ubiquitylated cargoes, as well as of Notch and Dome in endosomes. (A-F) High magnification of a region of mosaic eye imaginal discs (A-D), or of FE (E-F) shows accumulation of ubiquitylated cargoes in mutant cells (GFP-negative), as revealed by an antibody against mono- and polyubiquitin chains (Ubi). High magnification of the boxed areas is shown in insets.
(G-H) Mutant FE cells (GFP-negative) show accumulation of the Notch receptor. Notch receptor has been revealed using anti-NICD specific to the intracellular domain of Notch. Apical as well as intracellular accumulations of Notch ICD epitope is seen in *Hrs* and *Stam* FE mutant cells. High magnification of the boxed areas is shown in insets.
(I-I’) Co-localization with anti Notch ECD and Avl, marking early endosomes in mosaic eye imaginal discs. Notch is mainly accumulated in early endosome in GFP-negative mutant tissue.
(J-J’) Mosaic eye imaginal discs were stained with Ubi and anti-Domeless (Dome). *Hrs*, *Stam* mutant cells (GFP-negative) accumulate ubiquitylated cargoes and moderate levels of Dome, compared to WT.
(K-N) Endocytic trafficking assay with anti-Notch ECD to label Notch at the surface of living imaginal discs. In WT tissue, after labeling (0hrs), Notch is present mostly at the apical surface of the cell. After a 5-hour chase (5hrs) Notch is completely degraded in WT but still present in endosomes in *Stam* mutant discs, indicating that Notch is internalized but it is not degraded.

The Notch receptor is a cargo prominently subjected to endosomal sorting in *Drosophila* discs and FE cells [11–13]. To assess whether Notch is sorted and degraded in endosomes of ESCRT-0 mutant cells, we immunolocalized Notch in *Hrs*, *Stam* single or double *Hrs, Stam* mutant cells and in *Hrs Stam l(2)gl* triple mutant cells. Compared to WT cells, mutant eye disc cells displayed accumulation of Notch (Fig. 2G–I; Fig. 5G, I). Similarly, we found accumulation of Domeless (Dome), the single-pass non-tyrosine-kinase receptor for JAK/STAT signaling (Fig. 2J; Fig. 5H).

To follow sorting and degradation of transmembrane proteins over time, we performed a Notch endocytic trafficking assay in living imaginal discs [22]. Briefly, we cultured freshly dissected discs in insect media in presence of a Notch antibody that recognizes an extracellular epitope. We then washed and chased internalization of the bound antibody overtime. In contrast to WT discs, but like *Hrs* and *Vps25* mutant discs [22], *Stam* mutant, or *Hrs Stam l(2)gl* triple mutant disc cells displayed various degrees of intracellular signal after a 5 hrs chase, indicating that they fail to degrade endosomal Notch (Fig. 2K–N; Figure 5J–K). Co-staining of Notch with the early endosomal marker Avalanche (Avl) reveals that undegraded Notch and ubiquitin accumulate for the most part in early endosomes (Fig 2I; Fig. 5I). Overall, these data are consistent with a general defect in endosomal sorting and degradation of ubiquitylated cargoes, including signaling receptors, in ESCRT-0 mutants.

### ESCRT-0 is not required for endosome maturation

Given the inability of ESCRT-0 mutants to sort ubiquitylated receptors in endosomes, we next assayed whether mutant cells possess mature endosomes. One aspect of endosome maturation involves formation of ILVs during MVE biogenesis. To test whether ILV formation occurs in ESCRT-0 mutants, we analyzed the morphology of mutant cells at the ultrastructural level. To this end, we generated eye discs mutant for *Stam*, or *Hrs, Stam l(2)gl* or *Vps25,* encoding the eponymous ESCRT-II component. Compared to WT cells, *Vps25* mutant cells contain no MVE (Fig. 3A–B), as previously reported for *Drosophila* mutants in ESCRT-II components [18]. In these cells, we often observe the presence of very large clear vacuoles (Fig. 3B). Due to the loss of apico-basal polarity of *Vps25* mutant cells, we also find large interstitial spaces (Fig. 3B) In contrast, in tissue mutant for *Stam,* we find presence of normal MVEs (Fig. 3C). Similarly, in *Hrs, Stam, l(2)gl,* despite the presence of tissue disorganization similar to that of *Vps25* cells, due to the *l(2)gl* mutation, MVEs are present (Fig. 3D). These data are consistent with previous results in *Drosophila* Garland cells [23] and indicate that, different to ESCRT-II, ESCRT-0 components are dispensable for MVE biogenesis in epithelial tissue.

**Figure 3.**
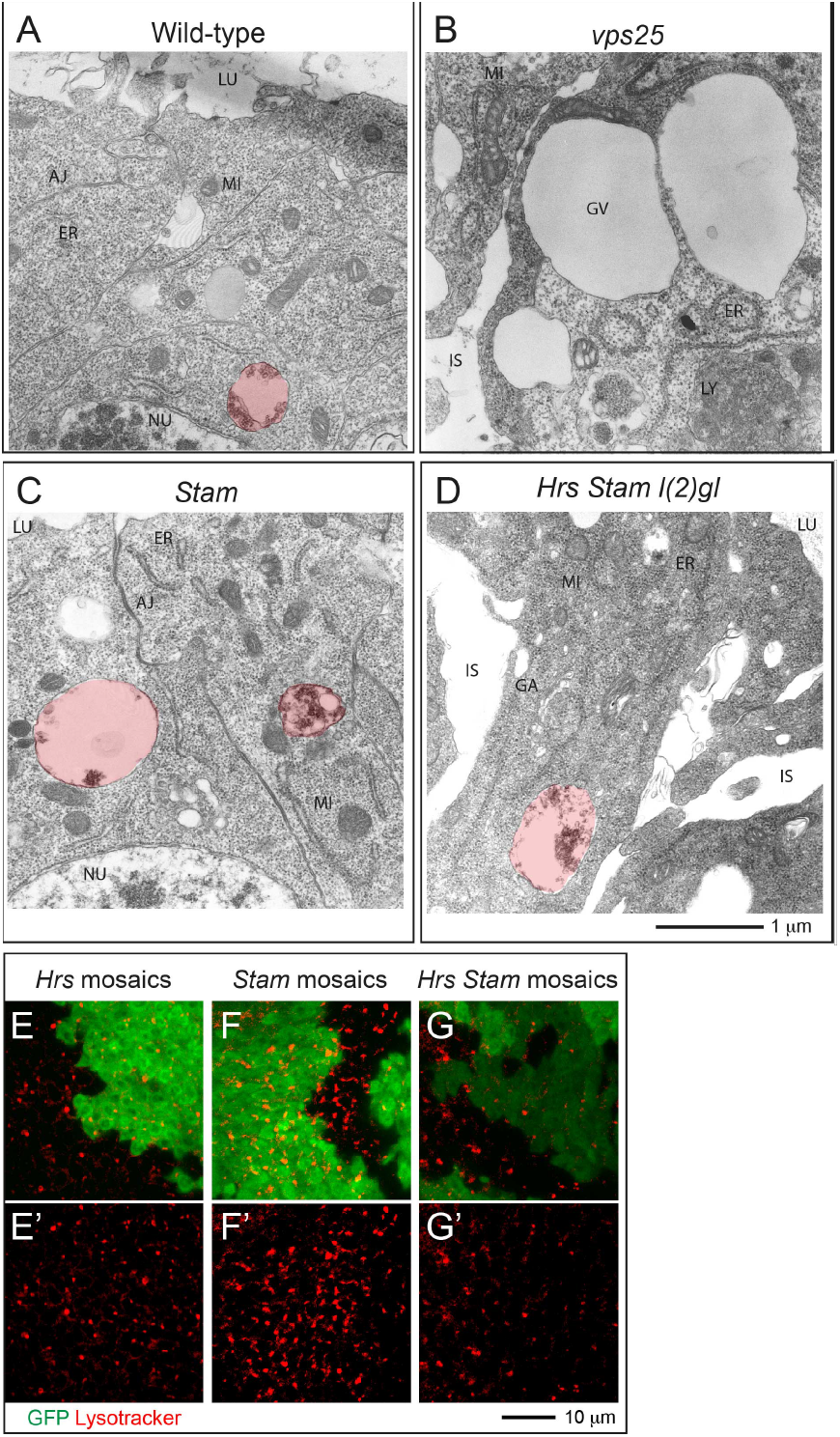
ESCRT-0 mutants do not affect endosomal maturation. (A-D) Electron micrograph of a section of eye disc tissue of the indicated genotype. A portion of the apical part of 2-3 epithelial cells above the level of the basal nuclei is shown. While MVEs (highlighted in red) are absent in *Vps25* mutant cells, they are present in ESCRT-0 mutant cells. Labels are as follows: LU: Apical lumen, AJ: Adherens Junctions, ER: Endoplasmic Reticulum, GA: Golgi apparatus, MI: Mitocondrium, NU: Nucleus, GV: giant vacuoles, IS: interstitial space between unpolarized cells.
(E-G) Incorporation of Lysotracker in mosaic discs. A single subapical confocal cross-section is shown in each panel, showing no difference in acidification in WT (GFP-positive) versus mutant cells.

Another aspect of endosomal maturation is the progressive acidification of the lumen of endosomes. To test whether *Hrs*, or *Stam* or *Hrs* and *Stam* mutant cells possess acidic endo-lysosomal organelles, we cultured mosaic discs in presence of Lysotraker, a vital dye that concentrates in acidic compartments. Compared with WT cells, clones of *Hrs* mutant cells incorporate equal levels of Lysotracker, consistent with previous evidence [31] (Fig. 3E). Similarly, *Stam* or *Hrs Stam* mutant cells are indistinguishable to surrounding WT cells, indicating no impairment of the ability to acidify endocytic organelles (Fig. 3F–G). Taken together, these data suggest that loss of Hrs, Stam or both do not affect endosomal maturation.

### ESCRT-0 is not required for Notch signaling activation or downregulation

Accumulation of Notch in endosomes of ESCRT-I, -II, -III mutants correlates with ectopic and ligand-independent Notch signaling [18]. In contrast, mutations that disrupt earlier steps of endocytic vesicle trafficking such as those affecting *Rab5* and *avl*, inhibit activation of Notch [22]. Therefore, it is unclear what to expect in ESCRT-0 mutants. To assay Notch activation in mutant FE cells, we monitored expression of the transcription factors Hindsight (Hnt) and Cut, whose expression is modulated by Notch activation at mid-oogenesis. In fact, upon expression of the ligand Delta in germline cells at stage 6 of oogenesis, Notch signaling is activated in neighboring FE cells. As a result, FE cells downregulate Cut expression, unpregulate Hnt expression, arrest mitotic cell cycles and begin to endoreplicate [32, 33]. Surprisingly, the pattern of Hnt and Cut expression detected by immunofluorescence in small clones of *Hrs,* or *Stam,* or *Hrs, Stam* mutant FE cells at stage 5-7 was unchanged, when compared to WT cells, indicating that Notch activation is not altered in ESCRT-0 mutants (Fig. 4A–F). In clear contrast, precocious expression of Hnt before stage 6 is observed in ESCRT-I mutant FE cells [22]. In *Hrs Stam l(2)gl* triple mutant cells, ectopic Hnt expression and failure to downregulate Cut expression is visible only in multilayering cells that do not contact the germline and are likely to not be reached by the ligand (Fig. 5L–M). Overall, our data suggest the ESCRT-0 activity is not required for Notch signaling and that its loss do not promote ectopic, ligand-independent activation, as observed in other ESCRT-I, -II, -III mutants.

**Figure 4.**
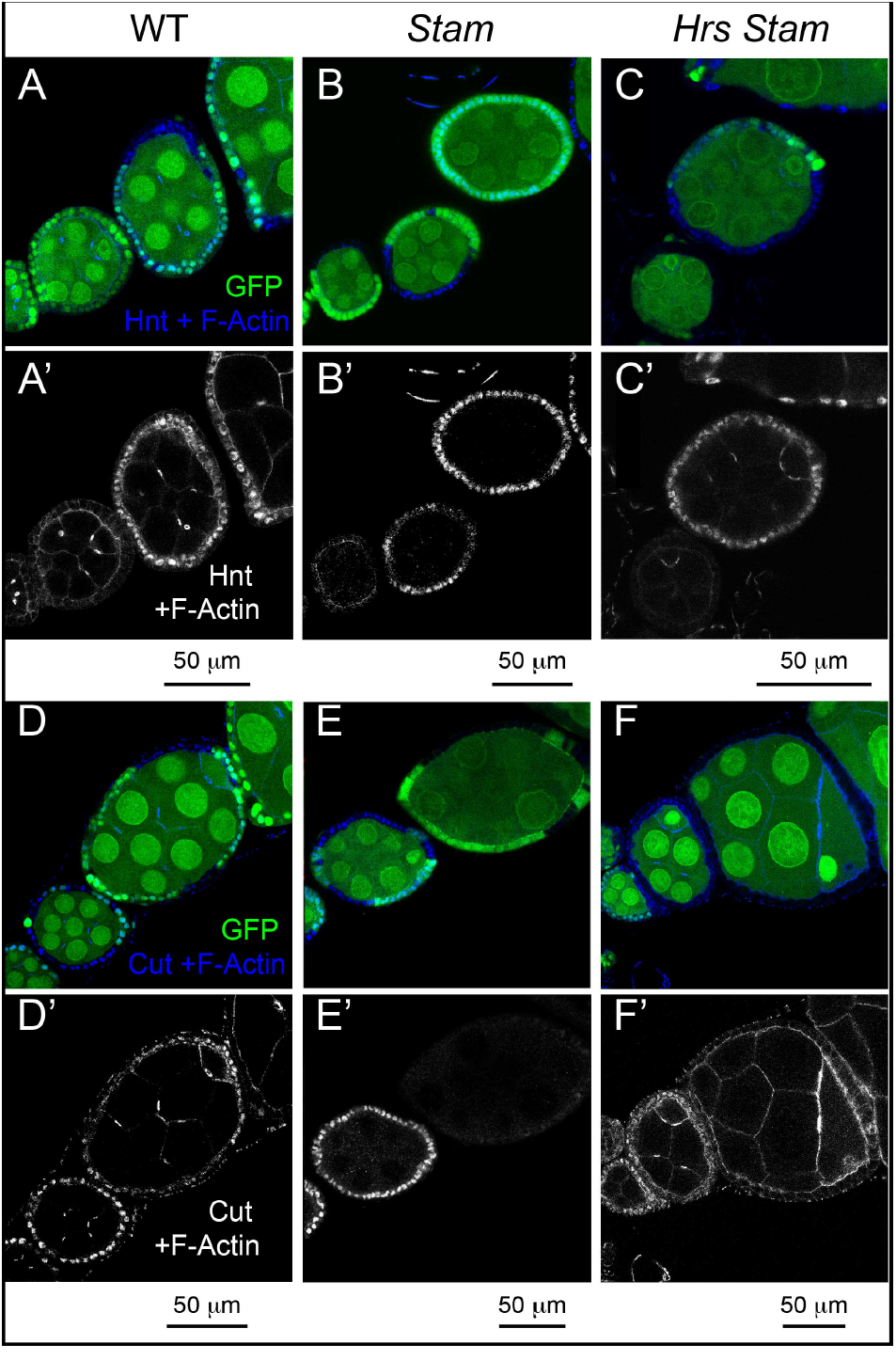
ESCRT-0 mutations do not alter Notch signaling in FE cells. Mosaic egg chambers at stages 5-7 of oogenesis stained to detect the Notch targets Hnt (A-C) and Cut (D-F) and f-Actin. *Stam* or *Hrs Stam* mutant cells are marked by the absence of GFP. In both *Stam* and *Hrs Stam* mutant FE cells, Hnt is normally expressed and Cut normally downregulated after stage 6, indicating no impairment of Notch signaling activation. (A’-F’) show single channels.

## Discussion

### The ESCRT-0 complex is dispensable for tumor suppression in *Drosophila*

In this study, we study the effects of impairment of ESCRT-0 function on *Drosophila* epithelial tissue development *in vivo*. We found that the recently reported [23, 24] *Hrs^D28^ Stam^2L2896^* double mutant allele carries a third mutation in *l(2)gl,* which we show is responsible for the loss of tumor suppressor (TS) phenotype of triple mutant tissue (Fig. 5). We analyzed independent *Hrs^D28^ Stam^2L2896^* recombinants devoid of *l(2)gl* mutations and observed that, when in homozygosity, these do not possess ability to growth into neoplasms, indicating that ESCRT-0 function *per se* is not tumor suppressive in *Drosophila*. While it is not clear when the reported *Hrs^D28^ Stam^2L2896^* chromosome [23] acquired the previously unrecognized *l(2)gl* mutation, it is possible that some of the phenotypes reported in the literature are due to impairment of *l(2)gl* activity.

**Figure 5.**
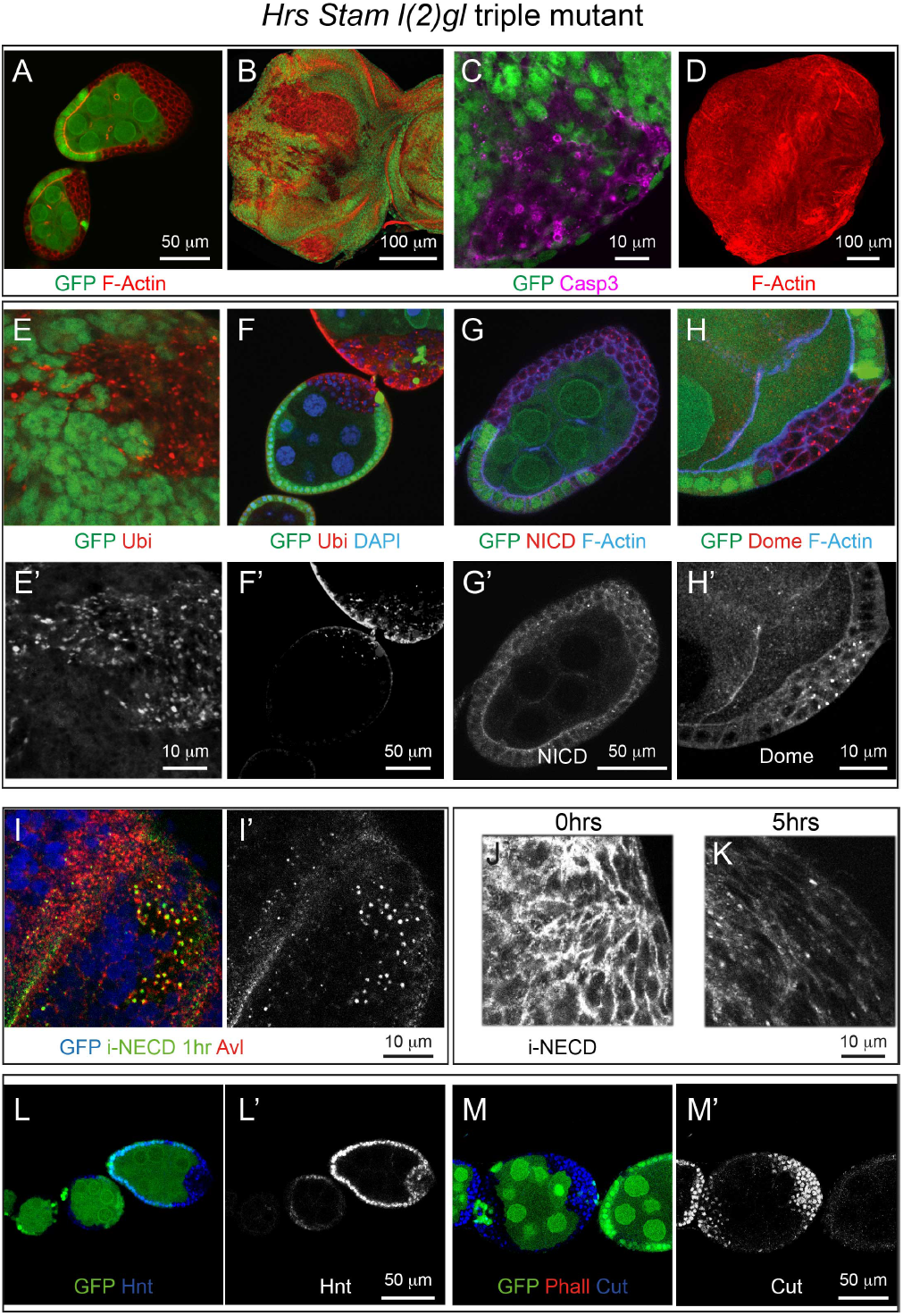
*l(2)gl* is responsible for the loss of tumor suppression phenotype in triple mutants. (A-B) Epithelial disorganization in *Hrs, Stam l(2)gl* triple mutant tissues revealed by staining to detect sub-cortical f-Actin. FE cells homozygous for *Hrs, Stam l(2)gl* (marked by the absence of GFP) in a stage 5-6 egg chamber are misshapen and multilayered. Eye imaginal cells homozygous for the mutations also show mesenchymal-like cells and autonomous disruption of epithelial organization.
(C) Mosaic eye imaginal discs stained with antibody anti-activated Caspase 3 to mark apoptotic cells. In *Hrs Stam l(2)gl* mutant tissue (GFP negative) apoptosis is activated.
(D) Eye imaginal disc formed by mutant cells homozygous for *Hrs, Stam l(2)gl* stained to detect sub-cortical f-Actin show a tumor-like phenotype.
(E-H) *Hrs Stam l(2)gl* mosaic eye and FE cells (marked by the absence of GFP) stained to detect ubiquitin, Notch and the Domeless receptors. Separate channels are shown in E’F’G’H’. Cells homozygous for *Hrs Stam l(2)gl* show accumulation of ubiquitin, Notch and Domeless intracellularly.
(I-I’) Co-staining of Avl with internalized anti Notch ECD (iNECD 1hr) in mosaic eye imaginal discs mutant for *Hrs Stam l(2)gl* (GFP-negative) reveals that undegraded Notch receptor is accumulated in Avl-positive endosomes.
(J-K) Endocytic trafficking assay with anti-Notch ECD to label Notch at the surface of living *Hrs Stam l(2)gl* mutant eye disc. Notch receptor fails to be degraded in mutant cells and it remains accumulated intracellularly after 5 hrs after the endo of labeling (0hrs).
(L-M) Mosaic egg chambers at stages 5-7 of oogenesis stained for Hnt and Cut. *Hrs Stam l(2)gl* homozygous cells are marked by the absence of GFP. In *Hrs, Stam l(2)gl* triple mutant cells Hnt expression and failure to downregulate Cut expression is visible only in multilayering cells that are likely not reached by the ligand, indicating that Notch activation is not affected in mutant cells that are exposed to the ligand and Notch is not ectopically activated in those that are not. WT controls for all panels are presented in Fig.1–2, 4.

The fact that ESCRT-0 is dispensable for tumor suppression (TS) marks a striking difference to mutations in most genes encoding components of the downstream ESCRT-I, -II, -III complexes. The discrepancy between the tissue architecture phenotypes of ESCRT-0 and other ESCRTs could be explained by different scenarios that we discuss below.

1. ESCRT TS function is not linked to endosomal sorting: ESCRT-III is very ancient, it is present in archaea and unicellular organisms [34], in which its membrane bending capacity is mostly used in the last step of cytokinesis [35, 36]. In contrast, ESCRT-0 has evolved recently, is dispensable for completion of cytokinesis, and might represent a specialization to sort a subset of cargoes in endosomes [37]. However, we do not favor the idea that the tumor suppression activity of ESCRT complexes correlates with their involvement in cytokinesis. In fact, while cell division and cytokinesis defect have been extensively linked to tumorigenesis, ESCRT-II, another ESCRT complex that behaves as TS, is dispensable for cytokinesis [35, 36].
2. ESCRT TS function is linked to endosomal sorting and residual Hrs or Stam function might be present in mutants: We think this is unlikely because both *Hrs^D28^* and *Stam^2L2896^* are null alleles to the best of our knowledge. In fact, *Hrs^D28^* expresses only the amino terminal first quarter of the protein, and is devoid of most functional domains [20], while *Stam^2L2896^* line harbors a non sense mutation leading to an early stop codon at amino acid 6 [24]. Both genes have no paralogs in *Drosophila*. In addition, quantitative RT-PCR also shows that in both *Hrs* and *Hrs Stam l(2)gl* mutants only 50% of the *Hrs* transcript is present. In both *Stam* and *Hrs Stam l(2)gl* mutant tissues, only 20-30% of the *Stam* transcript is present (Fig. 6A–C), indicating that in either backgrounds both mutant *Hrs* and *Stam* transcripts are possibly subjected to non sense-mediated decay, and further decreasing the likelihood of residual function.
3. ESCRT TS function is linked to endosomal sorting, but the relevant cargoes do not require Hrs or Stam for their sorting: Several studies suggested that alternative ESCRT-0 proteins may work in parallel, or even instead of Hrs and Stam. Good candidates are two families of proteins, GGAs and Tom1 (target of Myb1), both present in *Drosophila,* that have similar characteristics to those of ESCRT-0 components. These in fact contain VHSs, Ubiquitin binding, and Clathrin binding domains typical of ESCRT-0 components, they recruit ubiquitylated proteins to endosomal membranes, and they interact with ESCRT-I and Clathrin [38–40]. Thus, ESCRT-0 complex could be dispensable for sorting of TS-relevant proteins. Interestingly, endocytosis of junctional adhesion proteins, such as E-Cadherin, directly regulates polarity in *Drosophila* epithelia [41]. Consistent with a minor role of ESCRT-0 in controlling polarity, a study showed that mutation in *Drosophila Hrs* does not affect the localization of DE-Cadherin [21]. In contrast, junctional adhesion proteins appear sensitive to function of more downstream ESCRTs. Indeed, ESCRT-I and -III have been shown to be required for degradation of adhesive molecules, such as Claudin-1, and for maintenance of polarity in vertebrate epithelial cells [42]. Thus, we predict that proteins that play a role in ensuring correct epithelial architecture and polarity, such as those involved in cell-cell adhesion, might not require ESCRT-0 for their sorting and degradation.

**Figure 6.**
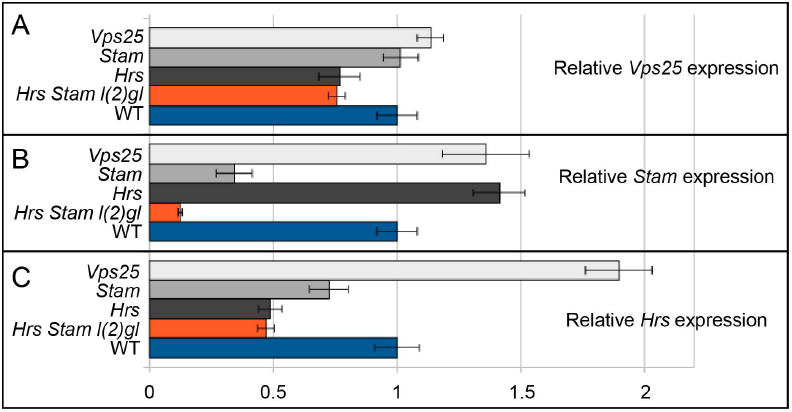
Mutant transcripts for Hrs and Stam are subjected to non sense-mediated decay. Quantitative RT-PCR experiment on mRNA extracted from eye imaginal discs from single Hrs or Stam or double Hrs, Stam or triple Hrs, Stam, l(2)gl mutant tissue compared to control indicates reduction of *Hrs* or *Stam* mRNA expression in corresponding mutant extracts.

### ESCRT-0 is dispensable for ectopic Notch activation in endosomes

Although in *Stam* and double mutants we observed accumulation of Notch receptor in endosomes, we were surprised to find no ectopic activation of Notch signaling. A trivial possibility to explain the difference is that Notch trafficking and degradation might be quantitatively more affected in ESCRT-I, -II, -III than in ESCRT-0 mutant, explaining the phenotypic differences. As discussed above, although our assays are not quantitative, the genetic nature of the mutants renders this possibility rather unlikely.

Another possibility is that recycling from endosomes to the plasma membrane might be important to prevent ectopic Notch receptor activation and recycling might still functioning in ESCRT-0 mutants, however it might not in ESCRT-I, -II, -III mutants. At present, whether the Notch receptor is recycled to the plasma membrane, and the status of recycling on different ESCRT mutant in *Drosophila* epithelia are unknown, preventing us to conclude on the likelihood of such an hypothesis. However, we observe an accumulation of Notch in endosomes of ESCRT-0 mutants that is comparable to that of ESCRT-I, -II, -III, an evidence that appears to contrast with the possibility of substantial recycling of Notch in ESCRT-0 mutants.

Finally, Notch accumulation in endosomes in ESCRT-0 mutants might not yield ectopic activation because such forced and ligand-independent Notch activation might require the cargo clustering by ESCRT-0 on the limiting membrane of endosomes. Alternatively, endosomes of ESCRT-0 mutant cells might not be mature enough to permit ligand-independent activation. These two not necessarily mutually exclusive possibilities are supported by the fact that Hrs and Stam act with Clathrin to trap and concentrate cargoes to be degraded on the endosomal membrane [9], and by evidence in *Drosophila* suggesting that ligand-independent Notch activation occurs in late endosome/lysosomes and depends on endosome acidification and maturation [31, 43–46]. Our data clearly indicate that two aspects of endosomal maturation, MVE biogenesis and endolysosomal acidification occur normally in ESCRT-0 mutants, suggesting that these could support later events required for ectopic Notch activation. The fact that ectopic Notch activation is not observed in the mutants thus points to cargo clustering as a potential prerequisite for ectopic activation of Notch. Whether cargo clustering is required for efficient Notch cleavage requires further studies.

In summary, our comparative analysis of *Hrs* and *Stam* in epithelial tissue *in vivo* reveals unexpectedly that ESCRT-0 is dispensable for control of cell polarity and proliferation, a major tumor suppressive event. Our data, marking a striking difference with ESCRT-I, -II, -III components, which act as TS, predict that specific cargoes important for cell polarity are sorted in endosomes independent of ESCRT-0 function, and that Notch activation, on the contrary, might be highly sensitive to receptor clustering by ESCRT-0 on the endosomal membrane.

## Acknowledgements

We thank Stephane Noselli, the BDSC and the DHSB for reagents. This work was supported by AIRC (Associazione Italiana Ricerca sul Cancro) strartup grant# 6118 (TV), by Telethon Italia grant# GGP13225 (TV), by Compagnia San Paolo (Torino) grant# 2011.1172 (CT), and by AIRC IG grant# 12035 (CT).

